# High-throughput method rapidly characterizes hundreds of novel antibiotic resistance mutations

**DOI:** 10.1101/2024.07.19.604246

**Authors:** Matthew J. Jago, Jake K. Soley, Stepan Denisov, Calum J. Walsh, Danna R. Gifford, Benjamin P. Howden, Mato Lagator

## Abstract

A fundamental obstacle to tackling the antimicrobial resistance crsisis is identifying mutations that lead to resistance in a given genomic background and environment. We present a high-throughput technique – Quantitative Mutational Scan Sequencing (QMS-Seq) – that enables quantitative comparison of which genes are under antibiotic selection and captures how genetic background influences resistance evolution. We compared four *E. coli* strains exposed to ciprofloxacin, cycloserine, or nitrofurantoin and identified 975 resistance mutations, many in genes and regulatory regions not previously associated with resistance. QMS-Seq revealed that multi-drug and antibiotic-specific resistance are acquired through categorically different types of mutations, and that minor genotypic differences significantly influence evolutionary routes to resistance. By quantifying mutation frequency with single base pair resolution, QMS-Seq informs about the underlying mechanisms of resistance and identifies mutational hotspots within genes. Our method provides a way to rapidly screen for resistance mutations while assessing the impact of multiple confounding factors.

## Main

Antimicrobial resistance (AMR) is an ongoing global health crisis limiting our capacity to treat infections. AMR infections lead to five million deaths each year, and this burden is predicted to increase considerably^1^. AMR often arises through chromosomal mutations that alter susceptibility to one or more antibiotics^2^. Consequently, predicting drug susceptibility from genomic data is becoming vital to antimicrobial stewardship by informing and stratifying treatments^3^. However, the accuracy of such predictions is limited by our existing knowledge of which mutations contribute to AMR and through what mechanisms.

Mapping mutations to AMR phenotypes is typically achieved through sequencing, either of clinical isolates or laboratory strains that have undergone selection for resistance. These approaches have proven powerful tools but are often biased towards mutations that confer high-level resistance, and struggle to discern how AMR evolution varies in different contexts. Smaller-effect mutations, epistatic interactions with genetic background, and environmental factors contribute significantly to overall resistance level, complicating genomic predictions^4^. As such, a key unmet challenge is to quantify the potential mutation space for resistance beyond the well-trodden evolutionary paths we currently understand.

To achieve this, we developed Quantitative Mutational Scan-Sequencing (QMS-Seq), an experimental evolution technique that adapts metagenomic sequencing to quickly characterize mutational landscapes for resistance. In QMS-Seq, random mutations are allowed to occur in the absence of selection, using subpopulations to limit founder effect bias and maximize diversity (**Fig. 1A**). Pooling the subpopulations creates a heterogeneous mixture mostly containing variants with only a single mutation. This population is spread across many selective plates – here containing the minimum inhibitory concentration (MIC) of an antibiotic. After resistant colonies have grown, they are mixed and sequenced collectively with sufficient depth to detect low-frequency resistance mutations. A custom bioinformatic pipeline (**Fig. 1B**) stringently filters for high-quality reads and utilises high specificity/sensitivity software^5^ to call single-nucleotide variants and small indels, while another tool, *breseq*^6^, identifies larger mobilization events for known insertion sequences. QMS-Seq’s experimental simplicity makes it ideal for high throughput methodologies, enabling quantification and combinatorial comparison across environmental conditions and genetic backgrounds.

**Fig. 1.**
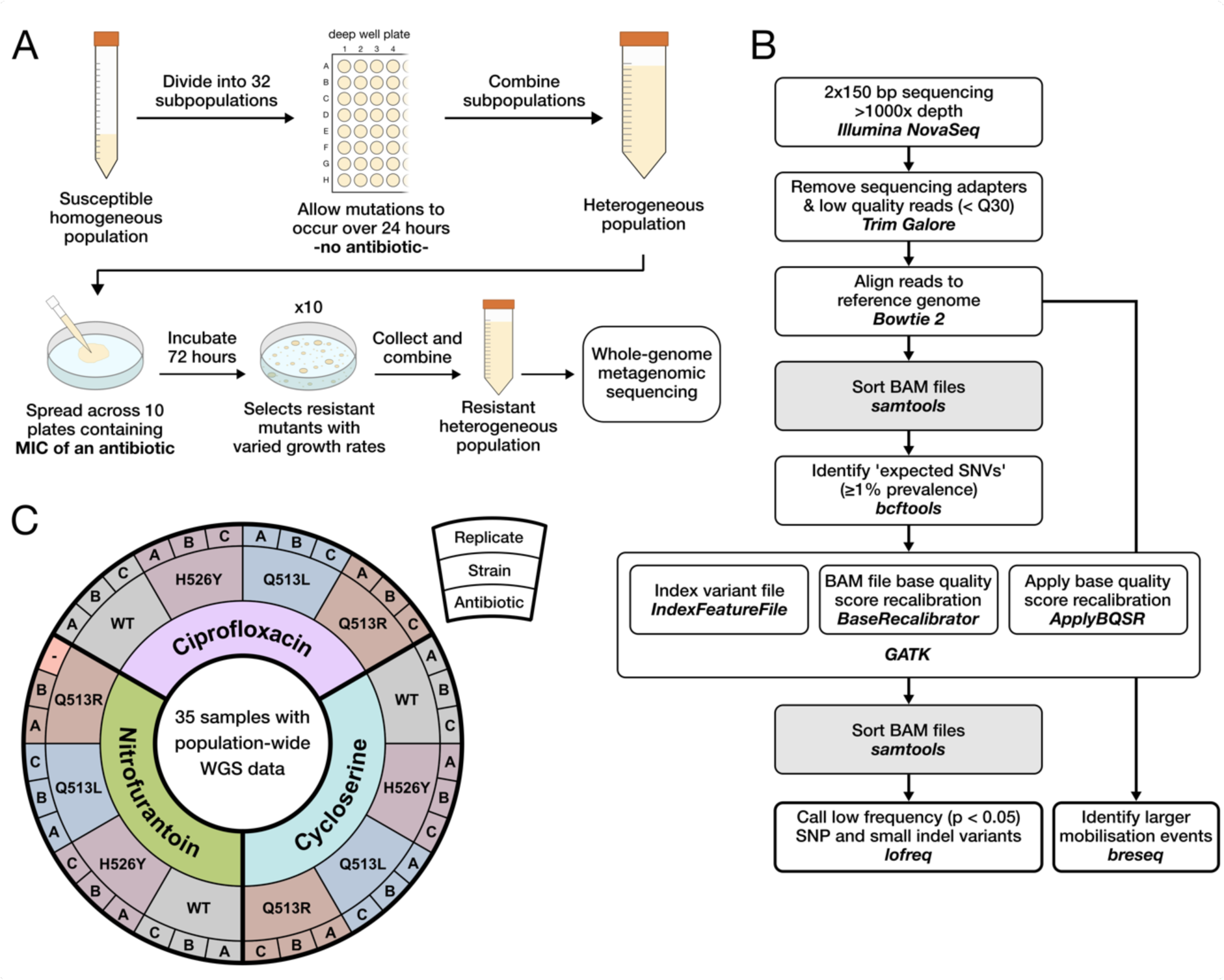
Quantitative Mutational Scan-Sequencing, QMS-Seq. (**A**) Experimental technique. (**B**) Sequencing and bioinformatics pipeline. (**C**) Experimental design. We tested three antibiotics and four strains of *E. coli*, performing three replicates for each of the 12 conditions, comprising 36 independent samples. The wildtype (WT) strain is BW25113, which served as ancestor for the three mutant strains. These each carry a different mutation in the *rpoB* gene which provides resistance to rifampicin and significantly changes their transcriptome compared to WT^7^. One nitrofurantoin/Q513R replicate was excluded from the analysis due to sequencing failure.

We employed QMS-Seq to investigate the mutational landscapes of four *Escherichia coli* strains for resistance to the MIC of three antibiotics with different mechanisms of action (**Fig. 1C**). This revealed many previously unknown resistance mutations, discriminated between mutations providing multi-drug or antibiotic-specific resistance, offered insights into the mechanisms underpinning resistance, and quantified how antibiotic class and genetic background influence AMR evolution.

## Results

### Mutational landscape for resistance

We observed 975 unique mutations across all conditions, in 278 genes and 62 regulatory features (**Fig. 2A, Supplementary Data 1-2**). We quantified the mutation *occurrence* – the number of independent samples each unique mutation was identified in. Interestingly, many more mutations occurred within intergenic regions (43%) than expected given the composition of *E. coli*’s genome (13% intergenic), suggesting that regulatory changes contribute to resistance evolution more than typically thought^8^ (**Fig. 2B**). Nearly half the mutations were in genes that have not previously been associated with resistance to any antibiotic, including 12 of the 30 most targeted genes. This indicates how much of the mutational landscape for resistance remains uncharacterized and emphasizes QMS-Seq’s utility in exploring it. Surprisingly, 15% of mutations affected genes known to be involved in antibiotic persistence. Persistence is a transient phenomenon that arises through the dysregulation of such genes^9^, yet we found heritable genetic changes, suggesting overlapping mechanisms for resistance and persistence at MIC. To verify the mutations were also relevant to non-laboratory settings, we assessed their prevalence in 21,711 clinical and environmentally isolated *E. coli* genomes from the BV-BRC database^10^ (**Fig. 2C**). Matching variants were found for 68 genes (of 278), with some identified in thousands of isolates including ciprofloxacin’s target *gyrA* and the three genes with highest mutation occurrence in our study (*ykfM*, *aceF,* and *ydbA*).

**Fig. 2.**
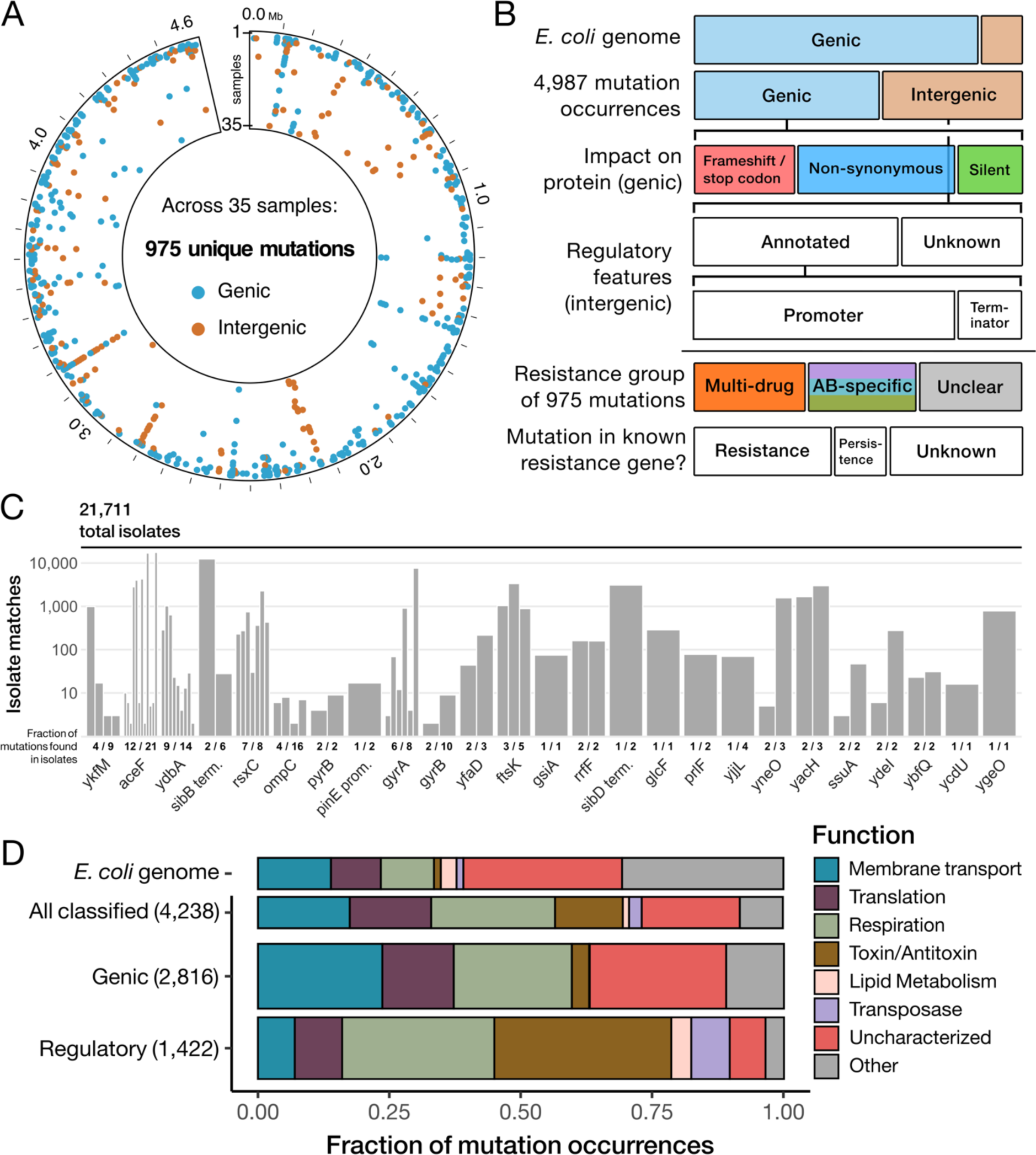
Summary of resistance mutations identified by QMS-Seq. (**A**) Circumferential axis is the mutation’s location in the *E. coli* genome (4.6 x 10^6^ bp). Radial axis is the mutation occurrence – the number of samples each mutation occurred in (out of 35). Colour indicates if the mutation occurred in a coding sequence (genic) or not (intergenic). (**B**) Genic mutations were further categorized based on how they affect the amino acid sequence; intergenic mutations based on whether they appear in known regulatory features^16–18^. Mutations were considered ‘multi-drug’ if they occurred in samples from each antibiotic, and ‘antibiotic-specific’ if they were significantly over-represented in one (*see Methods: Categorizing antibiotic-specific and strain-specific targets*). For every gene, any previous association with antibiotic resistance or persistence was confirmed through a literature search. (**C**) The 25 genes with the most isolates possessing matching mutations from the BV-BRC^10^, a WGS database for global human/animal-sourced *E. coli* isolates. Genes are ordered by their mutation occurrence in our samples. Data for all 68 genes and 3 regulatory regions with isolate matches in **Supplementary Data 3**. (**D**) All genic and annotated-intergenic (regulatory) mutation occurrences categorized by the biological function of the gene they affect. Also shown is the same classification of the WT genome (4,553 genes).

Most of the genes with known functions were involved with three broad biological processes: membrane transport, translation, or cellular respiration (**Fig. 2D**), which are all associated with antibiotic resistance and/or persistence^11–14^. The considerable variety of genes in each category suggests altering membrane architecture or core cellular processes is a frequent strategy for acquiring MIC-level resistance. There were notable differences between genic and regulatory mutations – the former much more common in membrane transporters, the latter accounting for 90% of toxin-antitoxin mutations. Toxin-antitoxin systems are thought to be a key driver of persistence^15^, indicating that the regulatory mutations we observed may recapitulate differential expression to provide a phenotype that mimics persistence, while being heritable rather than transient. Additionally, we observed mutations in multiple promoters for lipid metabolism genes and different copies of the *insH1* transposase, without equivalent genic mutations. Despite substantial overlap of functional categories between genic and regulatory mutations, the disparities indicate which resistance mechanisms favour differential expression over protein modification.

### Multi-drug vs. antibiotic-specific resistance

QMS-Seq differentiates between multi-drug resistance (MDR) mutations that confer resistance to all three antibiotics and mutations that confer antibiotic-specific resistance (ASR). We found categorical differences in both their intragenic positioning and their typical impact on the encoded protein. MDR mutations almost exclusively clustered within a small region of the gene, even for genes with many unique mutations (**Fig. 3A**). Furthermore, mutations leading to MDR were less likely to be ‘high-impact’ (frameshift or nonsense) while ‘low-impact’ (synonymous) mutations were significantly more frequent in MDR (ξ_8_ = 608; P < 0.001) (**Fig. 3B**). Conversely, for most of the top ASR targets, we identified high-impact mutations across the entire length of the gene, indicating loss-of-function provided resistance. Supporting this conclusion, over 75% of the insertion element mobilizations that we detected localized within ASR genes **(Supplementary Data 4)**. MDR mutations occurred more frequently than ASR mutations and accounted for nearly all regulatory mutations (93%), implying that expression-mediated resistance is less dependent on the antibiotic’s mechanism of action (**Fig. 3C**). Therefore, at MIC, the most common mutations are active site modifications and regulatory changes that provide MDR, whereas ASR typically arises from high-impact mutations.

**Fig. 3.**
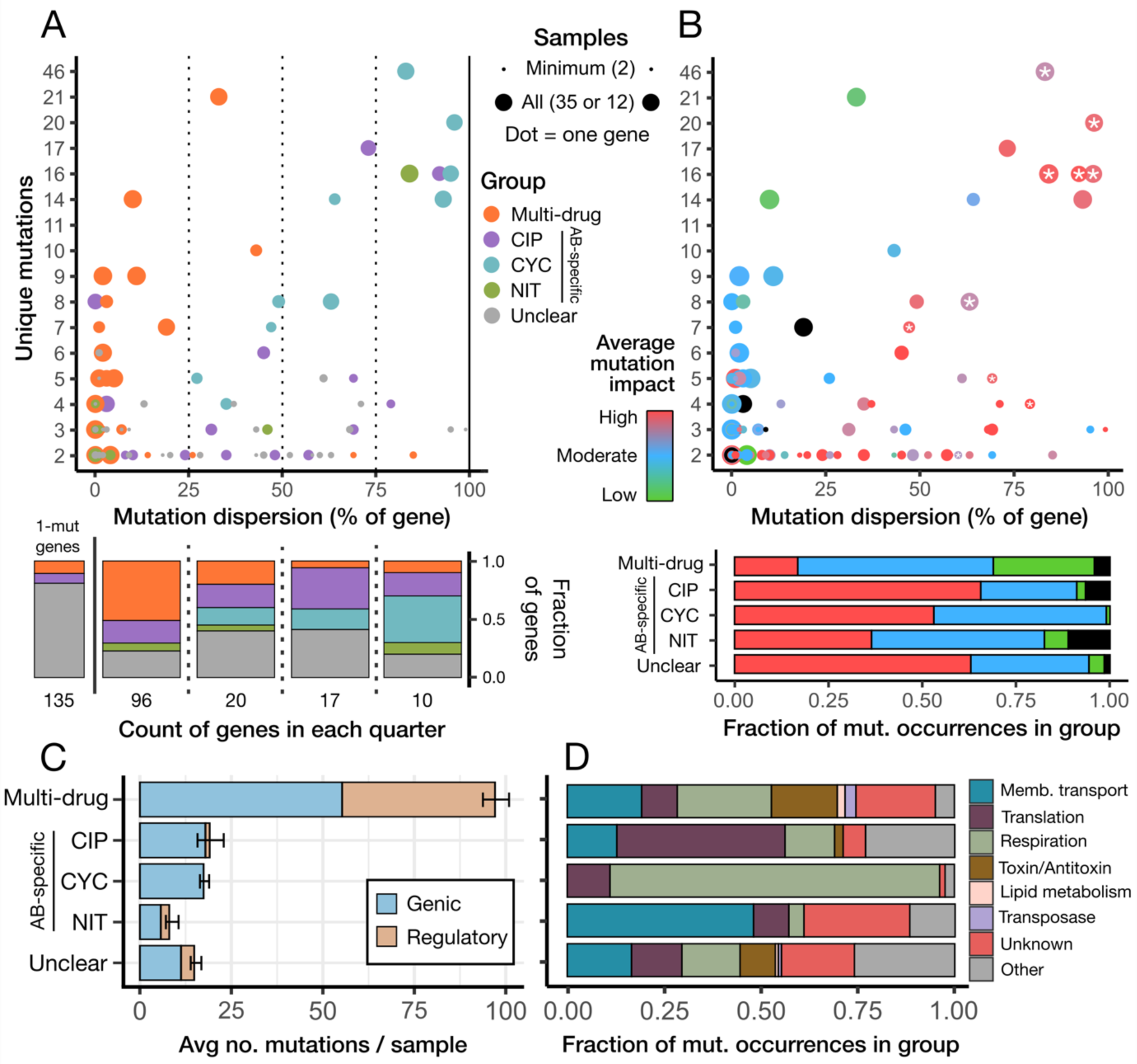
Multi-drug versus antibiotic-specific resistance mutations. (**A**) Every gene arranged by the number of unique mutations detected in that gene and the percentage of the gene’s length the mutations were dispersed across (*i.e.,* the distance between the furthest upstream and downstream mutations in the gene, see *Methods: Mutation impact and dispersion analysis*). Colour shows the resistance group classification of the gene. Size indicates the proportion of samples in which mutations were observed, out of a maximum of 12 for antibiotic-specific genes or 35 for multi-drug and unclear genes (those with insufficient mutation occurrences to confidently categorize). The bars below summarize each quarter of the plot, with an additional bar on the left representing 135 genes which cannot be displayed on the plot’s axes as only one unique mutation was observed. (**B**) The same plot, with points colored by the average impact-type of mutations observed in that gene. High: frameshift and nonsense, Moderate: in-frame insertion/deletion and non-synonymous, Low: synonymous. Black indicates a non-coding RNA. Stars indicate there was at least one occurrence of an insertion element mobilization event within the gene. Below is the impact-type distribution of all mutations occurring in each resistance group. (**C**) The mean number of mutations associated with each resistance group in any one sample. Error bars are SEM. (**D**) Proportion of mutations in each resistance group associated with different functional categories.

MDR mutations occurred in all the functional categories we identified, and almost every toxin-antitoxin mutation fell into this group (**Fig. 3D**). In contrast, ASR mutations were more likely to target specific functions, with translation favoured by ciprofloxacin (CIP), respiration by cycloserine (CYC), and membrane transport by nitrofurantoin (NIT). The absence of CYC-specific transporter mutations could be due to the drug’s disruption of cell wall biosynthesis, amplifying the fitness cost of mutations further affecting the membrane. As such, antibiotic mechanism of action may explain the observed biases in targeted biological systems, guiding the evolutionary pathways for ASR. Among the top ASR genes were well-documented targets of resistance mutations: *gyrA* and *icd* for CIP^19,20^, *ispA/B* and *ubiA/E/G/H* for CYC^21,22^, and *ompC* for NIT^23^. The latter, a membrane porin, is unique as it is typically considered a target for MDR, rather than being NIT-specific. However, despite mutations in every NIT sample, we identified none in CIP or CYC. Interestingly, with CIP’s other main resistance target (*gyrB*) we saw a cluster of CIP-specific mutations in the described quinolone-resistance region, but also several mutations outside this region in CYC and NIT (**Extended Data Fig. 1**), pointing to a potential role for *gyrB* in MDR.

### Genotype influences resistance evolution

We utilized four almost identical *E. coli* strains: the WT ancestor and three descendants, each with a single substitution in the *rpoB* gene encoding the β subunit of RNA polymerase **(Fig. 1C)**. The minor structural change provides resistance to rifampicin (which targets RNA polymerase) but also affects the transcription of many unrelated genes^7^. The different expression phenotypes associated with these mutations led us to hypothesize they may shape mutational pathways to secondary antibiotic resistance.

We quantified the differences in mutational landscapes associated with each strain, which showed a clear interaction between CIP and the Q513R strain – the number of unique genes targeted in this condition was significantly higher than any other (F_3,8_ = 6.2; P < 0.05) (**Fig. 4A**). CIP/Q513R mutation count was correspondingly high, on average nearly double the other conditions, which otherwise remained relatively consistent between antibiotics and strains (**Extended Data Fig. 2**). Over 70% of the mutations unique to CIP/Q513R were insertions or deletions, compared to ∼25% globally, suggesting a faulty DNA repair system played a role in the effect. In support of this, we identified several mutations exclusive to CIP/Q513R samples in critical genes for two DNA repair pathways, *recO* and *uvrB*^24,25^. Principal component analysis of the twelve conditions indicates the heavy influence this effect had on the mutational landscapes: while other conditions cluster by antibiotic, CIP/Q513R is responsible for most of the variance (**Fig. 4B**).

**Fig. 4.**
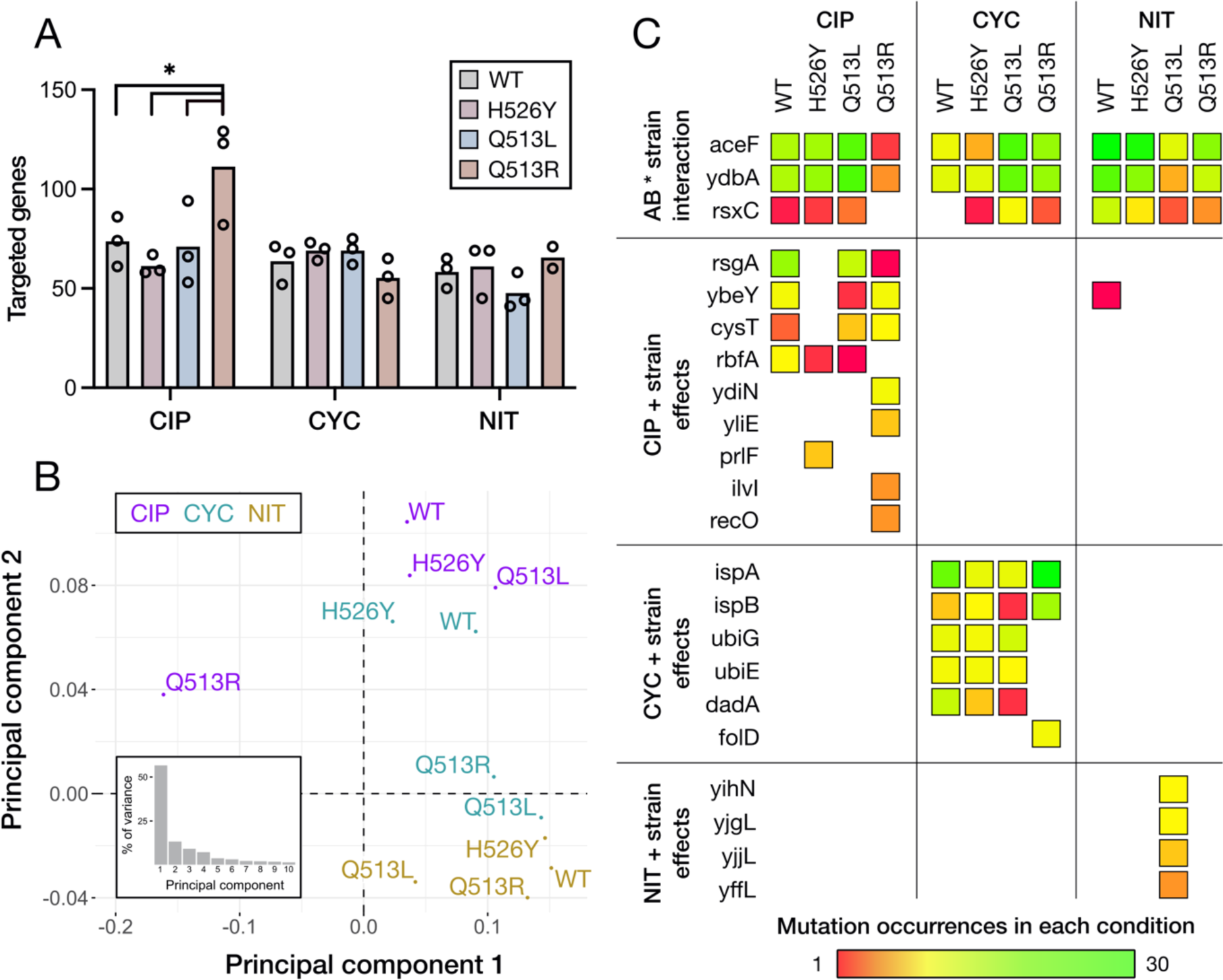
Influence of genetic background on resistance evolution. (**A**) The mean number of targeted genes (including regulatory targets) in samples from each of the 12 different conditions, with each individual replicate shown. (**B**) Principal component analysis of the 12 antibiotic/strain combinations, using the number of samples from each condition every mutation was observed in. Inset panel shows the scree plot indicating the percentage of variance described by each principal component. (**C**) The likelihood of a gene being targeted by mutations can depend on the strain’s genetic background. This was observed for MDR genes (top row) as well as ASR genes (bottom three rows). These are the top mutational targets with significant strain effects for each category, data for all genes in **Supplementary Data 2**.

At the individual gene level, strain-specific effects were more common in genes that were also specific to an antibiotic, although the mutational distributions of several MDR genes were shaped by an interaction *between* the antibiotics and strains (**Fig. 4C**). This was seen clearly in *rsxC*, which inhibits the SoxR MDR system partially responsible for clearing up reactive species^26^. Mutations were found across all antibiotics but were more common in NIT, which actively generates reactive species^27^. Curiously, WT was the most targeted strain in NIT but the least targeted across CIP and CYC, while the inverse was true for Q513L. Our data demonstrates how a single nucleotide polymorphism between strains can heavily influence evolutionary trajectories. While the exact relationship between the transcriptomic effects of rifampicin resistance mutations and secondary resistance remains of interest, QMS-Seq provides a way to evaluate the complicated interactions between antibiotics and genetic background.

### Case-studies demonstrate the depth of mechanistic insights afforded by QMS-Seq

#### Top genic target shows synonymous mutations can be under strong site-specific selection

The essential respiratory gene *aceF* is involved in catalysing pyruvate’s conversion to acetyl-CoA after glycolysis^28^. Previously found downregulated in persister populations^29^, it was the second highest target for genic mutations in our dataset. All the mutations were found in three homologous regions encoding lipoylation sites on the AceF protein^30^ (**Fig. 5A**). However, they were not distributed evenly, with only 2 mutation occurrences in site one compared to 64 in site two and 142 in site three – despite the repeat regions previously being considered redundant^31^. The functional site sequences are 71% identical, yet surprisingly *all* mutations occurred at positions that lack consensus between the three sites, invariably serving to bring one site into better alignment with another (**Fig. 5A**). Two-thirds of the mutations were synonymous, suggesting they affect translational efficiency, with the local translation rate potentially altering how the functional sites fold^32^. Irrespective of mechanism, the mutations in *aceF* exemplify the tight control of essential gene evolution, where marginal changes can give rise to important phenotypes such as antibiotic resistance. This analysis exhibits the resolution QMS-Seq can provide, offering mechanistic insights through the quantification of mutational landscapes, even at the level of individual genes.

**Fig. 5.**
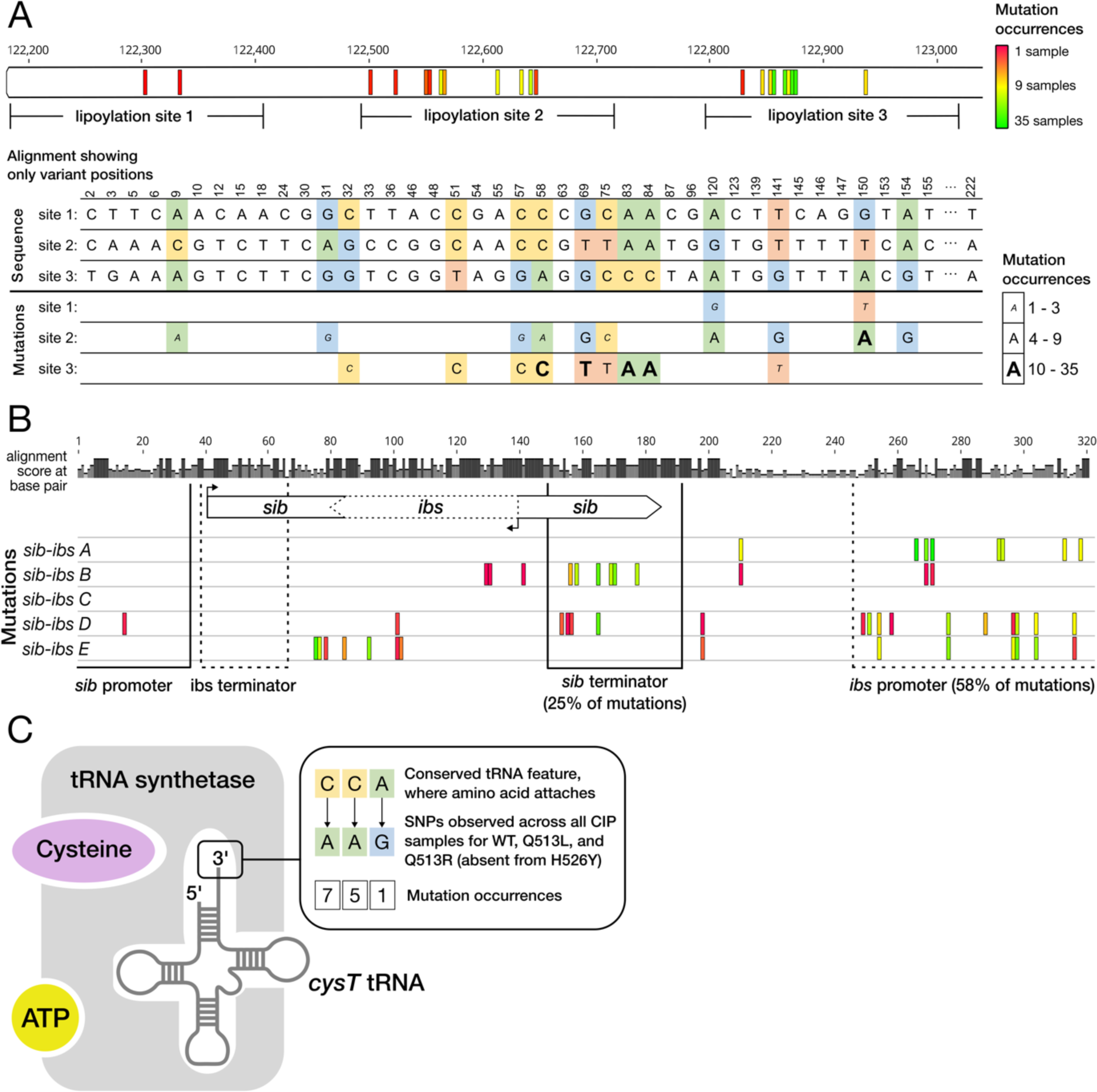
Case studies of notable resistance mutation targets. (**A**) The first 800 bp of the *aceF* gene (1,893 bp), a region encoding three homologous lipoylation sites. The 21 unique mutations are each colored by how many samples they occurred in. Below is a sequence alignment of the three functional sites, showing only the positions that are *not* conserved between all three sites (variant). Positions where a mutation occurred are highlighted, and the substituted base is shown below, indicating which site the mutation occurred in. 23 variant positions are not displayed because no mutations occurred nearby. (**B**) Sequence alignment of the five *sib-ibs* copies and their regulatory regions, with mutations colored by their occurrence as in (A). The alignment score indicates how many of the copies share consensus at each position. (**C**) tRNA aminoacylation (where *cysT* is charged with the cysteine amino acid by tRNA synthetase) and the mutations we observed in *cysT,* which in principle should prevent its interaction with cysteine.

#### Duplicates of a toxin-antitoxin system indicate the importance of regulatory mutations

The *sib-ibs* genes form a toxin-antitoxin system with an unclear physiological role^33^. The toxin and antitoxin have fully overlapping sequences but are transcribed from opposite strands, such that the longer antitoxin (a non-coding RNA) suppresses the complementary toxin mRNA. Five copies, in three locations, are present in the WT genome: *sib-ibs A/B* and *D/E* are found in tandem, while *sib-ibs C* is solitary. Fascinatingly, although the *A/B* and *D/E* regions were collectively the top mutational target, no mutations were observed in *sib-ibs C* despite significant homology with its cousins (**Fig. 5B**). Most of the mutations affecting *sib-ibs* copies occurred in annotated regulatory features, although the specific targets varied between copies. These mutations likely increase expression levels of the toxin, either directly or through inhibition of the antitoxin, aligning with previous reports that selective degradation of antitoxins facilitates growth arrest in persister populations^15^. Ultimately, the *sib-ibs* illustrate how different gene copies can have distinct evolutionary trajectories under antibiotic selection, either due to differences in sequence/function or their genomic location.

#### Should-be-lethal tRNA mutations provide ciprofloxacin resistance

We observed CIP-specific mutations in the *cysT* tRNA that altered its 3’ CCA tail, a conserved feature among tRNAs that is required for aminoacylation^34^ (**Fig. 5C**). Although unprecedented, this is not entirely without basis; co-treatment of CIP with cysteine (the amino acid for *cysT*) was shown to significantly increase killing of persister cells^35^. The synergistic relationship between CIP and cysteine appears to be recapitulated in our experiments, except through heritable mutations. However, *cysT* is the only cysteine tRNA in the genome, posing a puzzle as to how disruptions of its 3’ CCA are not lethal. *E. coli* carries a 3’ CCA repair mechanism^36^, which is likely able to rescue enough cysteine tRNA molecules to prevent death, presumably resulting in a variant with reduced translation of nearly every protein. Why this resistance strategy is unique to CIP, and no other tRNAs acquired genic mutations despite the abundance of translation-related targets, remains unclear. Furthermore, *cysT* is a prominent example of evolutionary differences between strains: all CIP samples contained at least one *cysT* mutation, except the three H526Y samples, which had none (**Fig. 4C**). Determining how *cysT* plays into the synergistic effect of CIP and cysteine will require further experimentation, but our findings highlight the new avenues for research opened by QMS-Seq’s ability to identify rare and low-fitness resistance mutations.

## Discussion

We used QMS-Seq to characterize the mutational landscapes for resistance to three different antibiotics, identifying nearly 1,000 mutations across over 300 coding and regulatory sequences. QMS-Seq provides a quantitative metric that captures how mutational effects depend on experimental conditions, enabling high-throughput assessment of the genetic and environmental factors that shape resistance evolution. Through this, QMS-Seq contributes novel insights into microbe biology while also addressing key deficiencies in the effort to monitor and track resistance evolution from genomic data. Its simplicity and broad applicability could help transition from a largely reactive response to the AMR crisis, towards pre-emptive strategies informed by studies examining many strains’ evolvability under diverse conditions.

QMS-Seq enabled the discovery of many genes and intergenic regions with previously unknown roles in resistance. This was facilitated by two design choices: (i) minimizing the capacity for high fitness variants to sweep through the population, therefore increasing diversity to capture a broader mutational landscape; and (ii) focusing on mutation *occurrence* by considering how often a mutation appears across independent samples instead of how prevalent a mutation is within a population. Furthermore, at MIC, relatively small phenotypic changes can provide resistance – in contrast to the majority of clinical isolate and experimental evolution studies that identify resistance mutations with populations exposed to much higher antibiotic concentrations. Using the MIC presumably contributed to the surprising abundance of regulatory and synonymous mutations, sites often thought to be under weak selection^8^. Such mutations may be enriched during selection at MIC because they are less likely to have a high fitness cost or be lethal^37^, and the effect they have on gene expression is sufficient to mediate low-level resistance. In known antibiotic persistence genes, which are canonically dysregulated to provide tolerance, regulatory or synonymous mutations may have conferred a permanent persister-like phenotype. The frequency of these mutations corroborates the notion that regulatory and synonymous mutations can significantly affect phenotype, particularly in environments where growth is restricted^38,39^.

The specific experimental design used with QMS-Seq can be tailored to answer a variety of questions about evolution. For example, using only one antibiotic and increasing the number of different strains could classify intraspecies variation in resistance evolution. This would help predict which lineages are most likely to acquire clinical resistance and how, a core goal of AMR surveillance. Variations to the method could also attempt to mitigate certain limitations of metagenomic sequencing. At present, QMS-Seq is unable to determine if secondary mutations have arisen after a resistance mutation, in the same lineage. Our filtering criteria (*see Methods: Filtering mutations under clear selective pressure*) verifies for strong selective pressure and should exclude hitchhiker mutations – but would not exclude subsequent mutations if they were compensatory and under strong positive selection in the given resistance background. This can be ameliorated by decreasing the growth period on selective agar (**Fig. 1A**), but this also reduces the capacity to detect rare and costly mutations.

QMS-Seq is experimentally simple, relatively inexpensive, and capable of rapidly identifying resistance mutations across an entire genome while distinguishing between those providing multi-drug vs. antibiotic-specific resistance. It affords sufficient resolution to build a map of which sequence features in those genes are targeted and how, helping to uncover the underlying mechanisms of resistance. Importantly, QMS-Seq evaluates the above in a strain-specific context, enabling researchers to disentangle how resistance evolution depends on the genotype of strains under investigation. In doing so, QMS-Seq can be employed to investigate questions critical to the assaying, monitoring, and stewardship of antimicrobial resistance, as well as provide a deeper view into the nature of microbial evolution.

## Methods

### Experimental strains

The ancestral WT was Escherichia *coli* BW25113 (NZ_CP009273.1) from the Keio collection^40^ containing a knock-out of the *tolC* gene. The three rifampicin-resistant strains were evolved from this ancestor. *rpoB* mutations were selected for as described in Soley *et al.*^7^. Whole genome sequencing was used to confirm that the strains differ by only one point mutation in *rpoB*. The *ΔtolC* strain was chosen so only *rpoB* mutations were selected for by rifampicin – TolC is a prominent efflux channel and common target for antibiotic resistance mutations. Its absence from the strains in this study may have increased the relative frequency of alternate resistance mutations and further contributed to the observed diversity. The three rifampicin-resistant strains each carried different amino acid substitutions: H526Y, Q513L or Q513R.

### Selecting for antibiotic resistance mutations

We inoculated Mueller-Hinton (MH) media with glycerol stock of each experimental strain, then split the inoculate into three tubes to start independent replicate populations. After overnight growth at 37°C we diluted the cultures 1:10,000 in MH and split them into 3 x 32 sub-populations in a deep 96-well plate. This was done to increase diversity and the relative frequency of low-fitness mutations by limiting founder effect bias or selective sweeps. We incubated the cultures for 24 hours at 37°C, allowing random mutations to occur during normal growth under minimal selective pressure. To select resistance mutations, the 3 x 32 subpopulations were then pooled into three independent heterogeneous populations and spread on selective plates containing the MIC of an antibiotic. We defined MIC as the lowest concentration of a given antibiotic that prevented any colonies from forming after three days of incubation. We determined the MIC of ciprofloxacin (0.005 μg/mL, Sigma-Aldrich), cycloserine (100 μg/mL, Sigma-Aldrich) and nitrofurantoin (10 μg/mL, TOKU-E) using MH agar plates. We determined the MIC for the WT strain and all three *rpoB* mutants separately, but no differences in MIC between strains were identified.

The three replicate populations were spread across 3 x 10 selective MH agar plates, using 200 μL of culture for each plate. The 30 plates were then incubated for three days to allow resistant mutants with severe fitness costs to form colonies. After three days, we collected and recombined colonies from the 10 plates into three heterogenous resistance populations to be sequenced. Prior to collection, we manually removed sections of the largest colonies, to increase the likelihood of identifying rarer and low-fitness mutations. QMS-Seq considers how many independent samples a mutation occurs in, rather than how prevalent it is within one sample population, so lowering the relative abundance of high-fitness mutants (which form large colonies) increases the relative abundance of low-fitness mutants, making them easier to detect without biasing later analysis.

To combine colonies from all 10 plates, fresh media was added to the plates and colonies were scraped to resuspend the cells, then tubes were thoroughly vortexed to break up any clumps. We measured optical density at 600 nm to estimate cell count and diluted the populations to 4.0 × 10^8^ cells/mL for storage at -80°C with 15% glycerol. The samples were later thawed, and genomic DNA was extracted using Genomic DNA Purification Kit (Promega). We verified the DNA concentration with a NanoDrop before sending for sequencing.

### Whole-genome metagenomic sequencing and low-frequency variant calling

We used Azenta Life Science’s Illumina NovaSeq 2×150 base pair sequencing package. They performed sequencing to a mean depth of at least 1000x per sample, a value estimated to allow detection of mutations with a frequency of at least 0.5% within the population. For QMS-Seq, raw sequencing reads were first trimmed with *Trim Galore* (v.0.6.7)^41^ to remove sequencing adapters and low-quality reads, with a Q score cut-off of Q30. Trimmed reads were aligned to the reference genome (*E. coli* K-12 BW25113) with *Bowtie2 (v.2.2.5)*^42^ using end-to-end alignment and the *–no-mixed* flag. BAM files were sorted with *samtools (v.1.16.1)*^43^, and single nucleotide variants (SNVs) with a minimum allele frequency of 1% were identified (these were considered ‘expected SNVs’) using the *bcftools (v.1.16) mpileup, call* and *view* commands. Prior to variant calling with *lofreq*, BAM files were pre-processed based on GATK best practice protocol as recommended by the *lofreq* documentation. Specifically, the variant file was indexed with *GATK (v.4.2.6.1)*^44^ *IndexFeatureFile* command and BAM files underwent base quality score recalibration with the *BaseRecalibrator* and *ApplyBQSR* commands, using the expected SNVs from the previous step as known sites. Recalibrated BAM files were sorted with *samtools*, and low frequency variants were called with *lofreq (v.2.1.3.1)* with a significance threshold of 0.05 and using the –*call-indels* flag.

We have taken several precautions to reduce the possibility of miscalls. We ensured a high-quality alignment with the use of paired end sequencing, stringent quality control of reads, and alignment of paired reads only, while excluding unpaired reads. *lofreq* itself utilizes mapping quality scores and base scores (which were recalibrated to reduce systematic errors in the scores assigned during sequencing) in calling low frequency variants. All samples had very high coverage and sequencing depth (**Supplementary Data 5**). We further checked the sequencing depth at the loci of the 5,037 filtered mutation occurrences – the median depth for all variant sites was 1151x (**Extended Data Fig. 3**).

The pipeline described above can identify SNVs and small insertions or deletions of less than 30 base pairs, at very low frequencies in the population. To identify low frequency mobilization events, which result in larger insertions and deletions, mutation calling was also performed using the *breseq* pipeline *(v0.38.1)*^6^. Trimming and read quality assessment was first performed on raw FASTQ files using *Cutadapt*^45^ and *FastQC*^46^ via the *Trim Galore* wrapper^41^, according to default run parameters. Mapping and variant calling was performed using *breseq* polymorphism mode with a “polymorphism-frequency-cutoff” of 0.001, with other run parameters set to default values. Reads were aligned to a modified version of the reference genome for which we have additionally annotated mobile elements^47^ as is required for the prediction of mobile element insertion events by *breseq*.

### Filtering mutations under clear selective pressure

Our variant calling pipeline identified 2,191 unique mutations across the 35 samples, which corresponded to 6,484 mutation occurrences, since many mutations were identified across multiple samples. The mutations were incredibly diverse, targeting over 900 different genes, around 1/5 of the WT genome. We were concerned that our pipeline, designed to pick up very low frequency variants in the population, had identified a host of additional mutations other than those providing antibiotic resistance. This seemed especially possible given the bacteria had three days of growth on the selective plate to accrue secondary mutations in addition to the resistance one.

To rectify this, we adopted a conservative filtering approach which identified true resistance mutations under clear selective pressure. Assuming a simplified possible mutation space of 1.4 × 10^7^ (the number of bases in the WT genome times three possible mutations at each base), and given an average of 185 mutations occurring per sample, the likelihood of the exact same mutation occurring in 2 or more samples (out of 35) by random chance is:

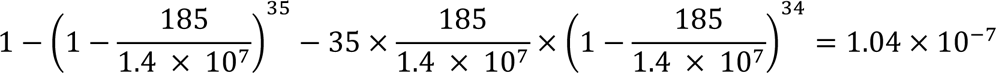

While the true possible mutation space is limited by essential bases and other factors, this number would not change by orders of magnitude. This calculation also excludes any other types of mutations, such as small insertions or deletions, and hence represents a conservative estimate. As such, we could be confident the 637 mutations which occurred in at least two samples were under strong positive selection.

This criterion was sufficient for intergenic and many genic mutations, however it excluded some genic mutations despite them being clustered in one gene. In certain genes, typically those targeted predominantly by nonsense or frameshift mutations, the *same* mutation would rarely appear in multiple samples, even if the gene was targeted by many *different* mutations across multiple samples. Hence, if there is insufficient pressure to mutate a specific base (or bases) in the gene over any other, the above criterion fails. To accommodate this, we also included 375 single-sample mutations observed in genes with more mutation occurrences than expected (**Extended Data Fig. 4).**

Our filtering criteria excluded 1,216 of the 2,191 mutations, which reduced the number of targeted genes from over 900 to 278. However, as our criteria generally filtered for mutations that occurred multiple times, the total number of occurrences only decreased from 6,484 to 4,987. This suggests that, while our filtering substantially decreased the observed variety of mutational targets, the filtered mutations still reflected most of the genetic diversity in the samples. To verify that the final 975 mutations we included in our analysis represented strong selective pressure, we compared their distribution throughout the genome to that observed by Foster *et al.* in a mutation accumulation study performed with *E. coli* in the absence of selection^48^. The gap sizes between mutations in our dataset indicated overwhelming clustering that was absent from the near-random distribution of 1,625 mutations observed by Foster *et al.* (**Extended Data Fig. 5**).

Finally, we estimated the typical number of mutations present in a single genome by performing WGS on 18 individual clones that had been streaked from the glycerol stock of nine different heterogeneous resistant population samples. Raw sequencing reads were first trimmed with *Trim Galore* with a Q score cut-off of Q20. Variants were then called with *snippy (v.4.6.0*). This confirmed that most clones did not contain more than one mutation (**Extended Data Table 1**).

### Annotating regulatory features in the genome

We mapped intergenic mutations to regulatory features using information from published databases: regulonDB^16^ and Ju *et al.*^17^, containing positions of known promoters and Rho-dependent/independent terminators in *E. coli*. An additional study was used to confirm the regulatory features of the *sib-ibs* genes^18^. In some instances, regulatory annotations overlapped with a coding sequence. Mutations that occurred in such regions were classified as regulatory if they were near (< 20 base pairs) the beginning or end of the gene.

### Classifying known resistance and persistence genes

We searched Google Scholar with the following queries: (“*gene name”* + antibiotic resistance + mutation) and (“*gene name*” + antibiotic persistence) for the 278 target genes. For 62 annotated regulatory regions, each gene in the transcriptional unit was inserted for *gene name*. If a publication was found that indicated mutations in the gene were associated with antibiotic resistance it was classed as such, regardless of the antibiotic or specific mutation(s). Antibiotic persistence genes were classified if they had specifically been implicated in persistence or found consistently dysregulated in persister populations. If no relevant publications appeared in the first 20 results, the gene was classed as not having been previously associated with antibiotic susceptibility.

### Categorizing the function of target genes

We used *EcoCyc*^49^ to annotate every target gene with a known biological role, including those immediately downstream of regulatory targets. To classify the genes into broad functional categories we used a combination of gene ontology (GO) biological process terms and published literature. For ‘membrane transport’, ‘translation’, ‘respiration’, ‘lipid metabolism’, and ‘transposase’ we identified the GO term(s) that were associated with each of these categories (**Supplementary Data 6**) and grouped target genes annotated with one of those terms. For transcription factors, which are annotated with GO terms indicating their enzymatic role rather than the downstream pathways they affect, classifications were supplemented with literature (**Supplementary Data 2**). This was also done for the ‘toxin-antitoxin’ category, as there is no relevant GO term. Target genes were classified as ‘other’ if they were annotated with GO biological process terms, but none fell under the above categories. They were classified as ‘uncharacterized’ if they were not annotated with any GO biological process, and we could not find a description of their function in literature. To construct an equivalent categorization of all the genes in the *E. coli* genome, we cross-referenced our list of GO terms against the GO annotations of the entire *E. coli* genome from *PANTHER*^50^, and identified the number of known toxin-antitoxin genes from literature^51^.

### Identifying matching mutations in global dataset of *Escherichia coli* isolates

To contextualize QMS-Seq mutations against global *E. coli* isolates, we downloaded all *E. coli* genomes from the BV-BRC database (as of April 2023)^10^. After filtering for high quality data from Illumina platforms and excluding isolates that lacked sufficient metadata, we kept 21,711 genomes for downstream analysis, consisting of 4918 raw sequence files and 16,793 assemblies. Variants were called using s*nippy* (*v4.6.0*) with the same reference genome used above (NZ_CP009273.1), and we counted the number of isolates with sequences matching mutations in our dataset.

### Categorizing antibiotic-specific and strain-specific targets

Given the quantity of mutations and their complex distribution across 12 experimental conditions, we chose to be conservative when categorizing mutational targets as ‘multi-drug resistance, MDR’ vs. ‘antibiotic-specific resistance, ASR’. For example, many mutations were identified in three different samples, two from one antibiotic and one from another. For mutations that appeared in only two samples from the same antibiotic, we could not be confident a hypothetical third occurrence would appear in the same antibiotic. Thus, we employed generalized linear modelling in *RStudio (v2023.09.1)* to statistically determine which genes/regulatory features were disproportionately targeted in one antibiotic. Targets with a mutation distribution where the antibiotic variable explained a significant (P < 0.05) amount of variation were classified as ASR. Targets that were mutated in at least one sample from *all* antibiotics were classified as MDR. Targets that were mutated in either one or two antibiotics, but with insufficient occurrences to pass our confidence threshold, were classified as ‘unclear’. The same approach was used to identify targets whose mutational distribution varied significantly between strains. We identified mutational distributions with higher-order interactions between antibiotic and strain, and targets where antibiotic and strain were both significant, but independent. We had the statistical power to identify targets with only strain effects (no antibiotic bias) but observed none. We used one-way ANOVA with multiple comparisons to examine differences in the total number of target genes (**Fig. 4A**) or mutations (**Extended Data Fig. 2**) between the 12 experimental conditions.

### Mutation impact and dispersion analysis

Predicting the impact (high, moderate, or low) of genic mutations on the encoded protein was automated by *SnpEff (v5.1)*^52^ which compares the variant sequences to the reference genome to predict the effect of single-nucleotide polymorphisms or small insertions/deletions. To calculate the average mutation impact for every targeted gene, each impact type was given a numerical value: high = 1, moderate = 0.5, low = 0. An average was taken for all the mutation occurrences in one gene to correspond with the three-color scale in **Fig. 3B**, red = 1, blue = 0.5, green = 0. We used Pearson’s Chi-squared test with Bonferroni correction and pairwise comparison to examine differences in the distribution of high/moderate/low impact types in MDR vs ASR mutations.

We complemented this analysis by calculating the ‘mutation dispersion’ for each gene, a metric quantifying how much of the gene’s total length mutations were identified across. Mutation dispersion was defined as the distance (in base pairs) between the furthest 5’ and furthest 3’ mutations, divided by the total length of the gene. When calculating mutation dispersion, we discounted ‘outlier’ mutations: low occurrence mutations located distantly from a cluster of high occurrence mutations. For each gene we considered every mutation occurrence as a number representing that mutation’s position between the first base of the gene and the last. We then calculated the interquartile range of these values and excluded mutations corresponding to outlier values. While still genuine resistance mutations, excluding the outlier mutations from this analysis provides a dispersion value that better reflects the way the gene is most commonly targeted.

### Principal component analysis of experimental conditions

We set up a table with 12 columns for each experimental condition and 975 rows for all identified mutations, then gave every mutation a value between 0 and 3 for each condition corresponding to the number of replicates the mutation was identified in. We used the *factoextra (v1.0.7)* package to perform principal component analysis and visualize the variables distributed along the first two components.

### Genome visualization and sequence alignment

We used *Geneious Prime® (v2023.3.2)* to visualize mutations throughout the reference genome in relation to the positions of coding and regulatory sequences. The annotated genome displaying the location and occurrence value of every mutation is available in **Supplementary Data 7**. This software was also used to perform *MUSCLE (v5.1)* sequence alignment for mutational targets with homologous regions, such as the *aceF* functional sites and *sib-ibs* copies.

## Acknowledgements

The authors thank Patricia Barkoci, Rebecca Boccola, Michael Brockhurst, Elizabeth Fozo, Rowan Green, Deniz Ozbilek, and Torsten Seeman. The authors acknowledge assistance from Research IT and the use of the Computational Shared Facility at the University of Manchester. Funding was provided by a Wellcome Trust and the Royal Society Sir Henry Dale Fellowship (grant 216779/Z/19/Z) and the EPSRC Centre for Doctoral Training in BioDesign Engineering (grant EP/S022856/1).

## Author contributions

Conceptualization: MJJ, JKS, SD, ML

Methodology: MJJ, JKS, CJW, DRG, ML

Investigation: MJJ, JKS, DRG

Visualization: MJJ

Funding acquisition: BPH, ML

Supervision: BPH, ML

Writing – original draft: MJJ

Writing – review & editing: MJJ, JKS, SD, CJW, DRG, ML

## Competing interests

Authors declare that they have no competing interests.

## Additional information

Supplementary Information is available for this paper.

Correspondence and requests for materials and data should be addressed to Mato Lagator.

## Data availability

Sequencing data has been deposited to NCBI [BioProject <u>PRJNA1071285</u>]. All processed and analysed data available upon request.

## Code availability

Scripts used to analyze sequencing data are available on GitHub [https://github.com/Lagator-Group/QMS-Seq/tree/main].

## Notes

### Competing Interest Statement

The authors have declared no competing interest.

