## Extended figures and table for "High-throughput method rapidly characterizes hundreds of novel antibiotic resistance mutations"

### Extended Data:

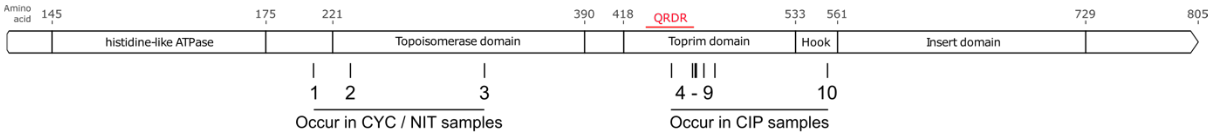


| Mutation information | | | | Number of replicates from each condition | | | | | | | | | | | |
|---|---|---|---|---|---|---|---|---|---|---|---|---|---|---|---|
| | | | | CIP | | | | CYC | | | | NIT | | | |
| Number | Reference | Alternative | Impact | WT | H526Y | Q513L | Q513R | WT | H526Y | Q513L | Q513R | WT | H526Y | Q513L | Q513R |
| 1 | T | G | Moderate | | | | | | | | | 1 | | | |
| 2 | G | T | Moderate | | | | | | 1 | | | | | | |
| 3 | A | C | Low | | | | | | | | 1 | | | 1 | 1 |
| 4 | T | G | Moderate | | | | 1 | | | | | | | | |
| 5 | G | A | Moderate | 3 | | | | | | | | | | | |
| 6 | C | T | Moderate | 1 | | | | | | | | | | | |
| 7 | T | G | Moderate | 1 | | | | | | | | | | | |
| 8 | G | C | Moderate | | | 1 | | | | | | | | | |
| 9 | TCACGACCGATAC | T | Moderate | | | 1 | | | | | | | | | |
| 10 | A | AT | High | | | | 1 | | | | | | | | |

**Extended Data Fig. 1. Mutations in *gyrB*.**

The GyrB amino acid sequence showing locations of known functional sites, and the positions of mutations observed by QMS-Seq. Most are mutations are specific to ciprofloxacin and cluster in or near the quinolone-resistance determining region (red). However, three other *gyrB* mutations occur in cycloserine and nitrofurantoin samples, in a different region of the gene.

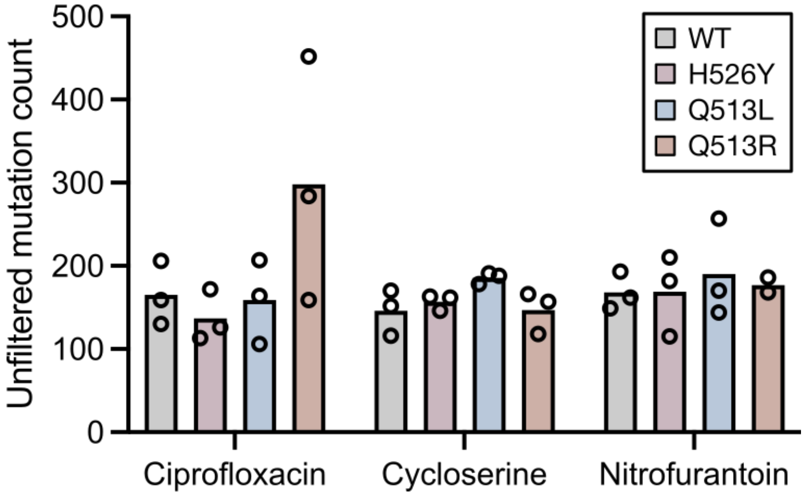


**Extended Data Fig. 2. Mutation count in every condition.**

The average number of unfiltered mutations identified in samples from the twelve different antibiotic / strain conditions. The value for each replicate sample is shown as a dot, bars are means.

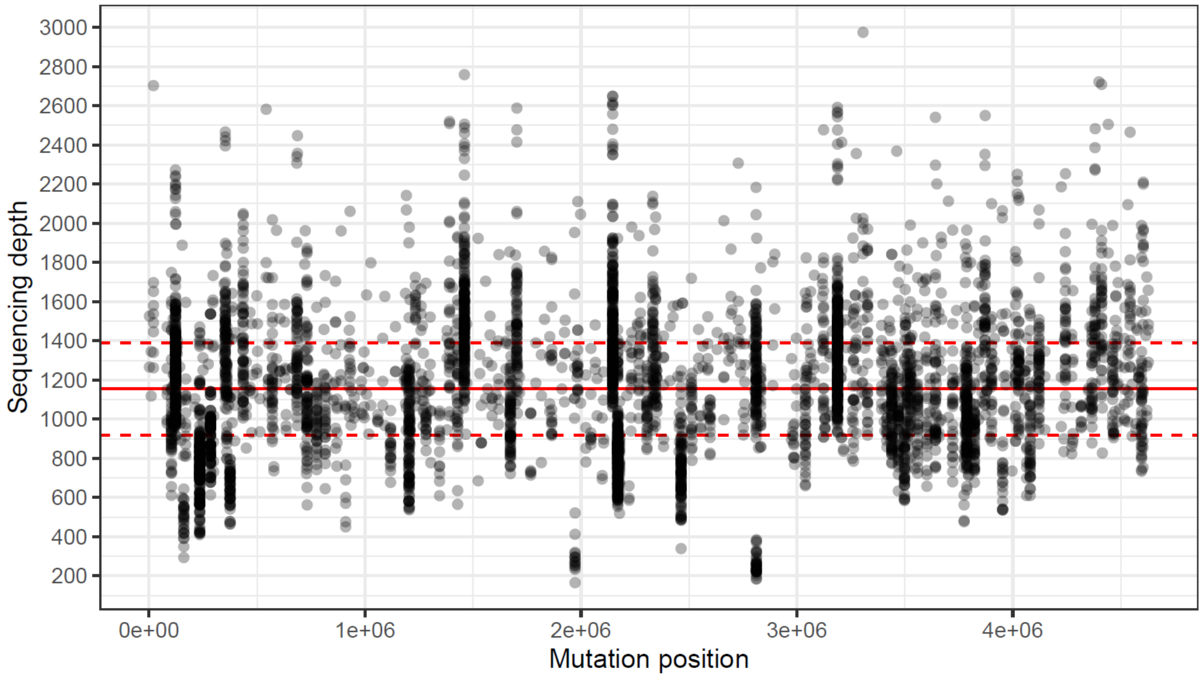


**Extended Data Fig. 3. Sequencing depth.**
Sequencing depth at the loci of each of the 4,987 mutation occurrences. Solid red line is the median, dashed lines are upper and lower quartiles.

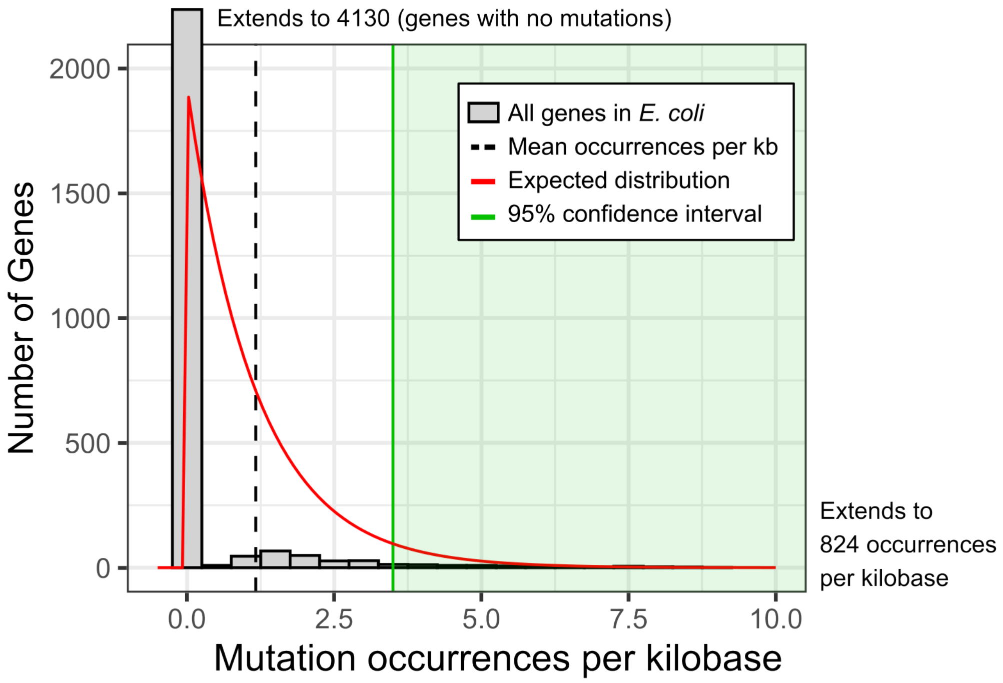


**Extended Data Fig. 4. Secondary filtering criterion.**
In addition to our primary filtering criterion (mutations must appear across two or more independent samples) we also included single-sample mutations in genes where significantly more mutations were observed than expected. To determine this, we calculated the number of mutation occurrences per kilobase for every gene in *E. coli*. We then included single-sample mutations occurring in genes with more mutation occurrences per kilobase than the 95% confidence interval, equivalent to > 3.5 mutation occurrences / kb.

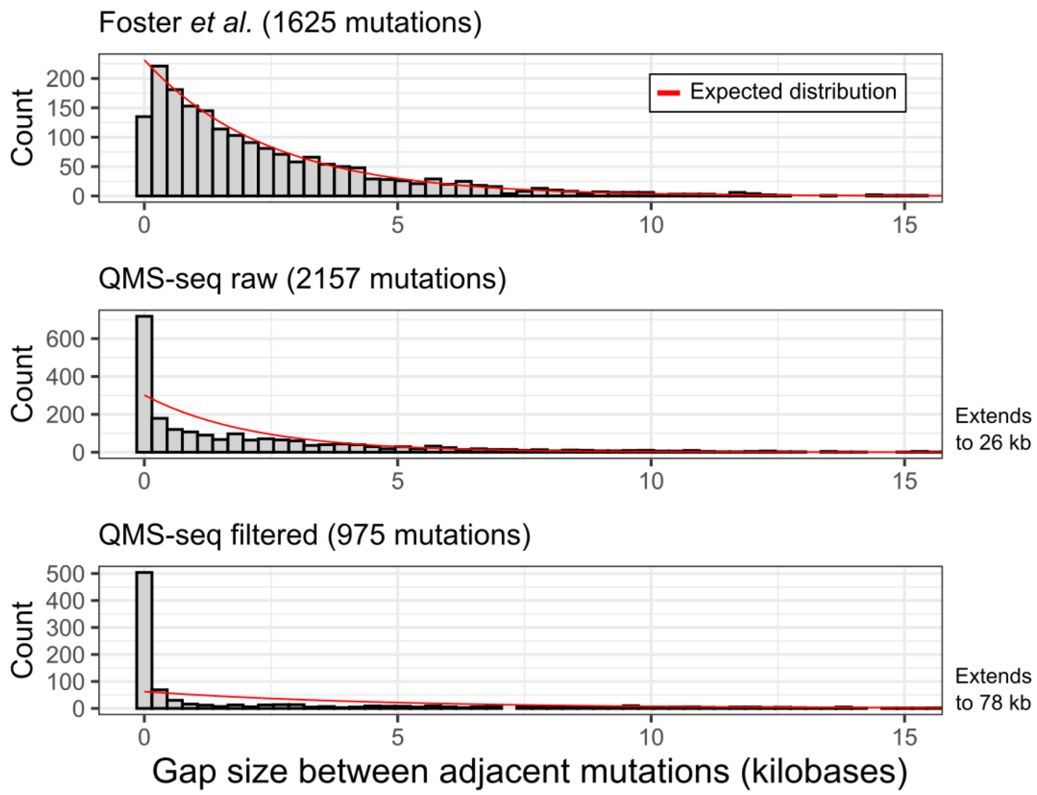


**Extended Data Fig. 5. Clustering of QMS-Seq mutations indicates strong selective pressure.**

Comparing the distribution of mutations throughout the genome between QMS-Seq and a mutation accumulation study performed in the absence of selective pressure using a Δ*mutL* strain of *E. coli* MG1655 (Foster *et al.* 2013)[48]. Absent from selection the mutations are spaced randomly, with the gap size between adjacent mutations nearly perfectly matching the expected exponential distribution. In contrast, the spacing of mutations identified by QMS-Seq shows significant clustering, indicative of selective pressure. The filtered QMS-Seq mutations (those analyzed in this study, *see Methods: filtering mutations under clear selective pressure*) show even greater clustering, suggesting the filtering has successfully discarded most hitchhiker mutations.

| Sample | Clone | Mutations |
|---|---|---|
| CIP/WT A | 1 | 1 |
| CIP/WT A | 2 | 1 |
| CIP/WT B | 3 | 1 |
| CIP/WT B | 4 | 0 |
| CIP/WT C | 5 | 0 |
| CIP/WT C | 6 | 1 |
| CIP/H526Y A | 7 | 1 |
| CIP/H526Y A | 8 | 0 |
| CIP/H526Y B | 9 | 0 |
| CIP/H526Y B | 10 | 0 |
| CIP/H526Y C | 11 | 2 |
| CIP/H526Y C | 12 | 1 |
| CIP/Q513R A | 13 | 1 |
| CIP/Q513R A | 14 | 0 |
| CIP/Q513R B | 15 | 1 |
| CIP/Q513R B | 16 | 1 |
| CIP/Q513R C | 17 | 1 |
| CIP/Q513R C | 18 | 0 |

**Extended Data Table 1. Validating the number of mutations per genome by whole-genome sequencing individual clones.**

We chose to examine CIP samples because it leads to DNA damage, and we surmised clones from these samples would have been most likely to acquire additional mutations. Most clones have one mutation per genome, only one was sequenced that had two. Several clones were sequenced with no mutation identified, these likely represent reversions of a high-fitness cost mutation, or cells which survived the antibiotic exposure via transient tolerance mechanisms. Reversions of structural variations (e.g. duplications or transpositions) are especially common, and both are major mechanisms of resistance evolution[53]. Clones with no mutation (and thus no accompanying fitness cost) would also be more prominent because of how we collected colonies to perform individual whole genome sequencing. We picked colonies that had grown over 16 hours after streaking from the resistant heterogeneous populations we sent for metagenomic sequencing. Clones in the heterogeneous population were grown on the plate for three days, as the growth rate of many colonies was very slow. This meant that, after 16 hours, only the most fit clones would be visible on the plate and then picked for sequencing.
